# The secret of VDAC isoforms is in their gene regulation? Characterization of human VDAC genes expression profile, promoter activity, and transcriptional regulators

**DOI:** 10.1101/2020.08.10.244327

**Authors:** Federica Zinghirino, Xena Giada Pappalardo, Angela Messina, Francesca Guarino, Vito De Pinto

## Abstract

**Background:** VDACs (Voltage-Dependent Anion-selective Channels) are pore-forming proteins of the outer mitochondrial membrane, whose permeability is primarily due to their presence. In higher eukaryotes three isoforms raised during the evolution: they have the same exon-intron organization and the proteins show the same channel-forming activity. We provide a comprehensive analysis of the three human VDAC genes (VDAC1–3), their expression profiles, promoter activity, and potential transcriptional regulators.

**Results:** VDAC isoforms are broadly but also specifically expressed in various human tissues at different levels with a predominance of VDAC1 and VDAC2 over VDAC3. However, RNA-seq CAGE approach revealed a higher level of transcription activation of VDAC3 gene. We experimentally confirmed this information by reporter assay of VDACs promoter activity. Transcription Factor Binding Sites (TFBSs) distribution in the promoters was investigated. The main regulators common to the three VDAC genes were identified as E2FF, NRF1, KLFS, EBOX transcription factors family members. All of them are involved in cell cycle and growth, proliferation, differentiation, apoptosis, and metabolism. More transcription factors specific for each isoform gene were identified, supporting the results in the literature, indicating a general role of VDAC1, as actor of apoptosis for VDAC2, and the involvement in sex determination and development of VDAC3.

**Conclusions:** For the first time, we propose a comparative analysis of human VDAC promoters to investigate their specific biological functions. Bioinformatics and experimental results confirm the essential role of VDAC protein family in mitochondrial functionality. Moreover, insights about a specialized function and different regulation mechanisms arise for the three isoforms genes.

## Background

VDAC (Voltage-Dependent Anion-selective Channel) is the prototype of the family of subcellular pores responsible for the permeability of mitochondrial outer membrane [1]. This protein is a key control point for the passage of ions and metabolites to guarantee cell energy production. During evolution more isoforms raised in the eukaryotic organisms [2]: they were characterized by the functional point of view, revealing very close permeability features [3–7]. Also, experiments aimed to study their expression patterns in different tissues indicated very subtle differences and a quite ubiquitous presence in the tested tissues [8].

In general, the distribution of VDAC isoforms is more or less ubiquitous in tissues but with a prevalence of VDAC1 isoform. Although the three VDAC genes codify proteins with apparently the same function, differences in the amino acid content, in the localization in the mitochondrial outer membrane layer [9], in the channel functionality and voltage dependence [6, 7], in the contribution of N-terminal portion to cell viability and survival [10], lead to hypothesize a more specialized role/function for each isoform in different biological contexts.

Results from various groups highlighted specific functions for each isoform, and it is considered a common notion that, for example, VDAC1 is a pro-apoptotic actor [11], while VDAC2 is an anti-apoptotic protein [12]. The role of VDAC3 was associated with sex tissue development and maintenance soon after its discovery [13, 14]. A further step was the deletion of single isoforms’ genes, which was performed in transgenic mice. In VDACs knock-out mice, physiological defects are linked to altered structure and functionality of mitochondria [11, 13]. Mitochondria lacking VDAC (VDAC1−/−), are characterized by increased size, irregular and compacted cristae, and altered respiratory complex activity particularly in various types of muscles [11]. Other alterations have been detected in specific tissue injuries or pathologies like Alzheimer’s disease [15, 16]. These abnormalities may concur to produce not functional cells. The absence of VDAC1 in muscle biopsies of child patients with impaired oxidative phosphorylation, suffering for mitochondrial encephalomyopathy, was associated with a fatal outcome [17, 18]. VDAC2−/− knock-out mice could not develop to adult individuals, but only ES cells could be grown [19]. A significant degree of growth retardation characterizes VDAC1/3 −/− mice; and VDAC3 −/− deficient male mice show the peculiarity to be infertile, with a disassembled sperm tail, the flagellum essential for sperm motility [13]. More recently the amino acid composition of the three mammalian VDAC isoforms was examined, showing interesting peculiarities [20]. The cysteine content of the three isoforms is different and this was correlated to their oxidation and potential disulfide bonds formation [21]. The three isoforms carry strikingly different amounts of cysteine whose different role has been hypothesized [20].

Similarly to VDACs knock-out mice, defects in the brain, muscle, and germinal tissues function were observed in *Drosophila melanogaster* porin1 mutants [22]. In the yeast *S. cerevisiae,* two *porin* genes were found but only *porin1* coding VDAC1 is essential for life [23].

A peculiar expression of VDAC isoforms was observed in cells and tissue of the germinal lines of different organisms. While VDAC1 is mainly located in cells of reproductive organs necessary to support the development of gametes [24, 25], VDAC2 and VDAC3 are expressed in a specific portion of sperm and oocyte and genetic variants or the aberrant regulation of these genes are correlated with infertility [26, 27].

Mechanisms of VDAC genes expression regulation have never been explored in a comparative and systematic way. Only one paper was published, describing the organization and activity of mouse VDAC gene promoters [28]. A prediction of the regulatory regions of the three VDAC isoforms show that the promoters lack the canonical TATA-box and are G+C-rich, the Transcription Start Site (TSS) was identified and the mouse putative VDAC promoters tested for their activity. In more recent years other few reports have been published on VDACs’ promoter regulation for specific germinal lineage. The activity of VDAC2 promoter in mammals developing ovary triggered by GATA1 and MYBL2 transcription factors lead to autophagy inhibition confirming its relevant role in cell survival [29]. In human male abnormal hypermethylation of VDAC2 promoter correlated with idiopathic asthenospermia while in complete unmethylation or mild hypermethylation, sperm motility improved confirming the role of VDAC2 expression in human spermatozoa [30]. The increased expression level of VDAC1 transcript was also induced by a lncRNA through enhancement of H3K4me3 levels in its promoter leading to apoptosis of placental trophoblast cells during early recurrent miscarriage [31]. For the first time, we propose with this work a study of the genomic region located upstream the TSS generating the three VDACs mRNA. We collected the information on human VDACs genes structures and transcription from the main available public resources. Once VDAC gene promoters were identified, we analyzed the sequences using bioinformatics software to identify Transcription Factors Binding Sites (TFBSs) distribution. We performed experimental tests with a molecular biology approach to confirm the insights obtained and to assess the transcriptional activity. We believe that understanding the molecular mechanisms triggering VDAC genes transcription in physiological and altered conditions might highlight the biological role of each isoforms inside the cells and in different biological contexts.

## Results

### Structure of human VDAC1, VDAC2 and, VDAC3 genes: transcripts variants and promoters

The vast amount of large-scale genomic projects, high-throughput sequencing, and transcriptomic data, as well as the plentiful supply of promoter resources, assist in the comprehensive reconstruction of a transcriptional regulatory region. To intersect these data enabled us to provide a framework of the main functional DNA elements for the identification of biochemical active regions supposed as gene promoter sequences of each VDAC isoforms, and to study the transcriptional control of the promoter structure by the analysis of TF-binding sites specificity and co-association patterns with other TFs.

Here, we will show the results of merging these clues with information from gene expression profile datasets, and with analysis of the promoter-specific activity of VDACs, which helped to investigate the co-expression relationship and the context-dependent regulation of VDAC genes.

Our approach narrowed down the analysis of the promoter region of each VDAC isoform to defined 600bp segment for P_*VDAC1*_ (chr5:133,340,230-133,340,830; hg19) P_*VDAC2*_ (chr10:76,970,184-76,970,784; hg19) and P_*VDAC3*_ (chr8:42,248,998-42,249,598; hg19), whose range is from −400 bp to +200 bp relative to TSS of annotated promoter sequences in EPD new (v.006). The boxed area in Figure 1 a-c highlights the UCSC-based BLAT result of P_*VDAC1*_, P_*VDAC2*_ and P_*VDAC3*_ matching with functional elements positioned within the region of accessible chromatin that define the nearest active promoter sequence. The structural organization of VDACs genes, transcripts, and the surrounding regions were analyzed here.

**Figure 1.**
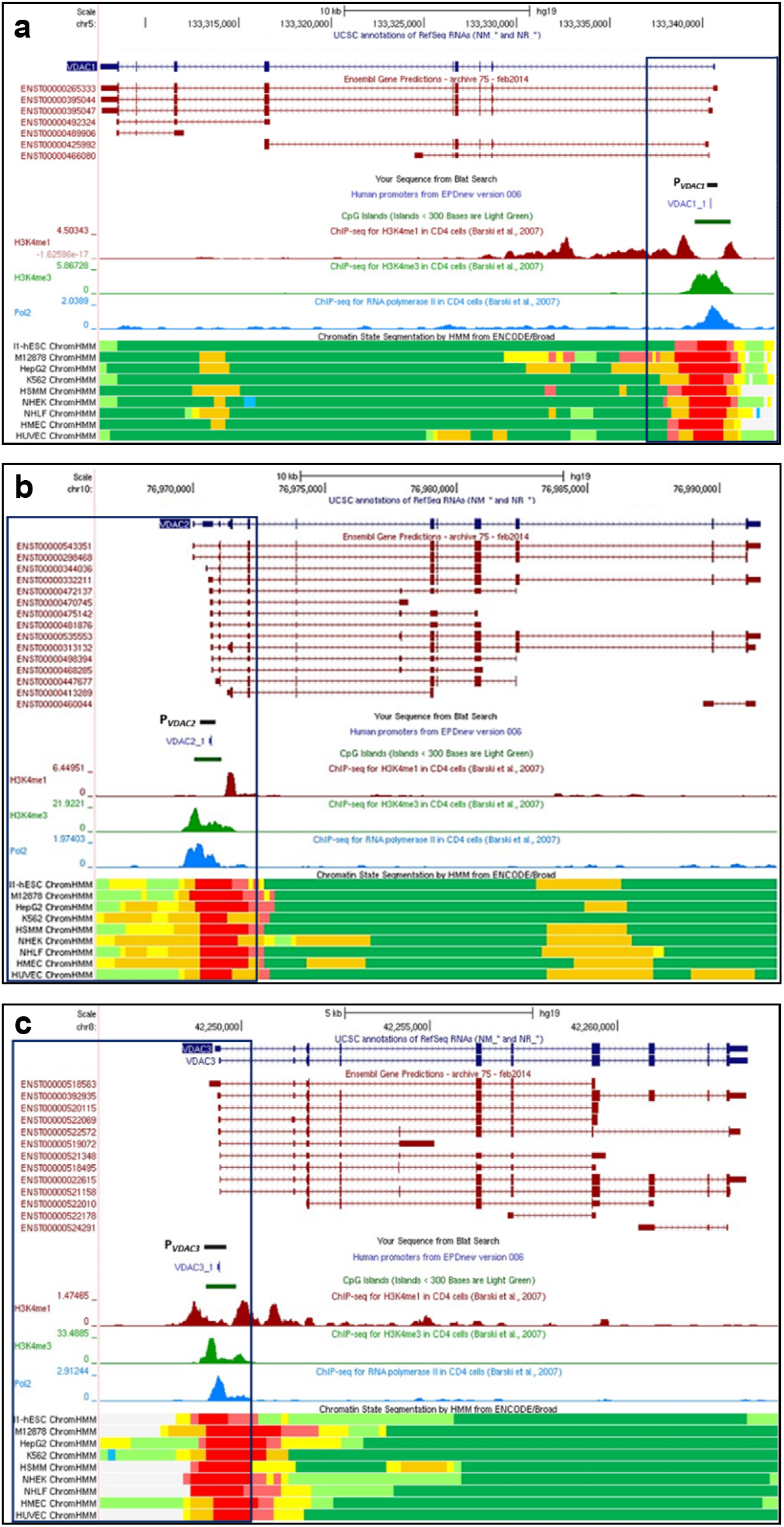
Human *VDAC* isoforms genes structure and function. Overview of gene structure of human VDAC isoforms and their most relevant functional and structural sites. (**a**) *hVDAC1* gene location on Chr5:133,340,230-133,340,830 (GRCh37/hg19) from UCSC Genome Browser; (**b**) *hVDAC2* gene location on Chr10:76,970,184-76,970,784; hg19; (c) *hVDAC3* gene location on Chr8:42,248,998-42,249,598; hg19. In each panel a black box encloses the 600bp promoter sequence indicated as P_VDAC1_, P_VDAC2_ and P_VDAC3_, respectively, and aligned with the annotated sequence from EPDnew (v.006). This allows highlighting the profile of transcriptional activity of the gene promoter region by CpG island identification, levels of enrichment of the H3K4me1 and H3K4me3 histone marks, and RNA Pol2 and Chromatin State Segmentation ChIP-seq data. Functional elements of Chromatin state segmentation by HMM of nine different cell lines are identified by different colours as follows: bright red: active promoter; light red: week promoter; orange: strong enhancer; yellow: weak/poised enhancer; blue: insulator; dark green: transcriptional transition/elongation; light green: transcriptional transcribed.

The transcripts coding the three VDACs functional proteins are reported with the code ENST00000265333.8 for VDAC1, ENST00000543351.5 for VDAC2, ENST00000022615.9 for VDAC3 corresponding respectively to NM_003374.2, NM_001324088.1 and NM_005662.7 in the Refseq database of NCBI.

Several transcript variants from Ensemble database are also indicated for each VDAC isoforms. Most of the splice variants have the same exons composition compared to the coding transcript but differ in the length of the 5’ and 3’. Some of them are processed transcripts, other features retained intron and for VDAC3 two are involved in nonsense-mediated decay mechanism. It is not known whether the other splice variants identified have any functional biological role, however, gene expression data collected from NIH Genotype-Tissue Expression project (GTEX), report the expression of them, including the non-protein coding transcripts.

The 600bp-selected promoter region shows a high degree of overlap with the identification of the CpG island and the RNA polymerase II binding site close to the TSS, also confirmed by ChIP-seq data of chromatin-state model and the enrichment levels of the H3K4me1 and H3K4me3 histone marks, chosen as the best predictors of transcription and open chromatin elements available among the UCSC regulation tracks.

### VDAC genes transcription by Gene Expression Atlas resources

Gene expression profile was highlighted through the analysis of available high-throughput data, included in international collaborative projects aimed to characterize human genome expression and regulation. In this work data, revised and curated by Gene Expression Atlas of EMBL-EBI, from GTEx [32] and RIKEN Functional ANnotation of the Mammalian Genome 5 (FANTOM) project [33], were reported for an interesting comparison of results obtained with RNA-seq methodology. The expression patterns of the three VDAC isoforms depict the level of VDAC transcripts distribution in different human tissues.

The RNA-Seq expression data from GTEx, obtained by human tissues samples from post-mortem individuals, show that the level of VDACs mRNA expression seems comparable among the three isoforms but with a prevalence of VDAC1 (Fig. 2 a-b). A relevant result is that all VDAC isoforms are mainly expressed in skeletal muscle and heart comparing the expression with the other tissues. VDAC1 and VDAC3 isoforms are expressed with a similar score while VDAC2 is expressed to a minor extent. Among the other tissues, we noticed that both VDAC1 and VDAC3 are represented in different portions of the brain with a higher score than VDAC2. However, the presence of VDAC1 and VDAC3 in brain regions seems to be differentiated since the former isoform is more expressed in diencephalon while VDAC3 in the telencephalon. A similar situation can be observed in organs forming the digestive apparatus. The specificity of VDAC isoforms expression can be highlighted in other tissues. For example, VDAC1 is revealed in skin tissues, exposed or not to sun, kidney, VDAC2 in the bladder, cervix/ectocervix, vagina, and VDAC3 in testis.

**Figure 2.**
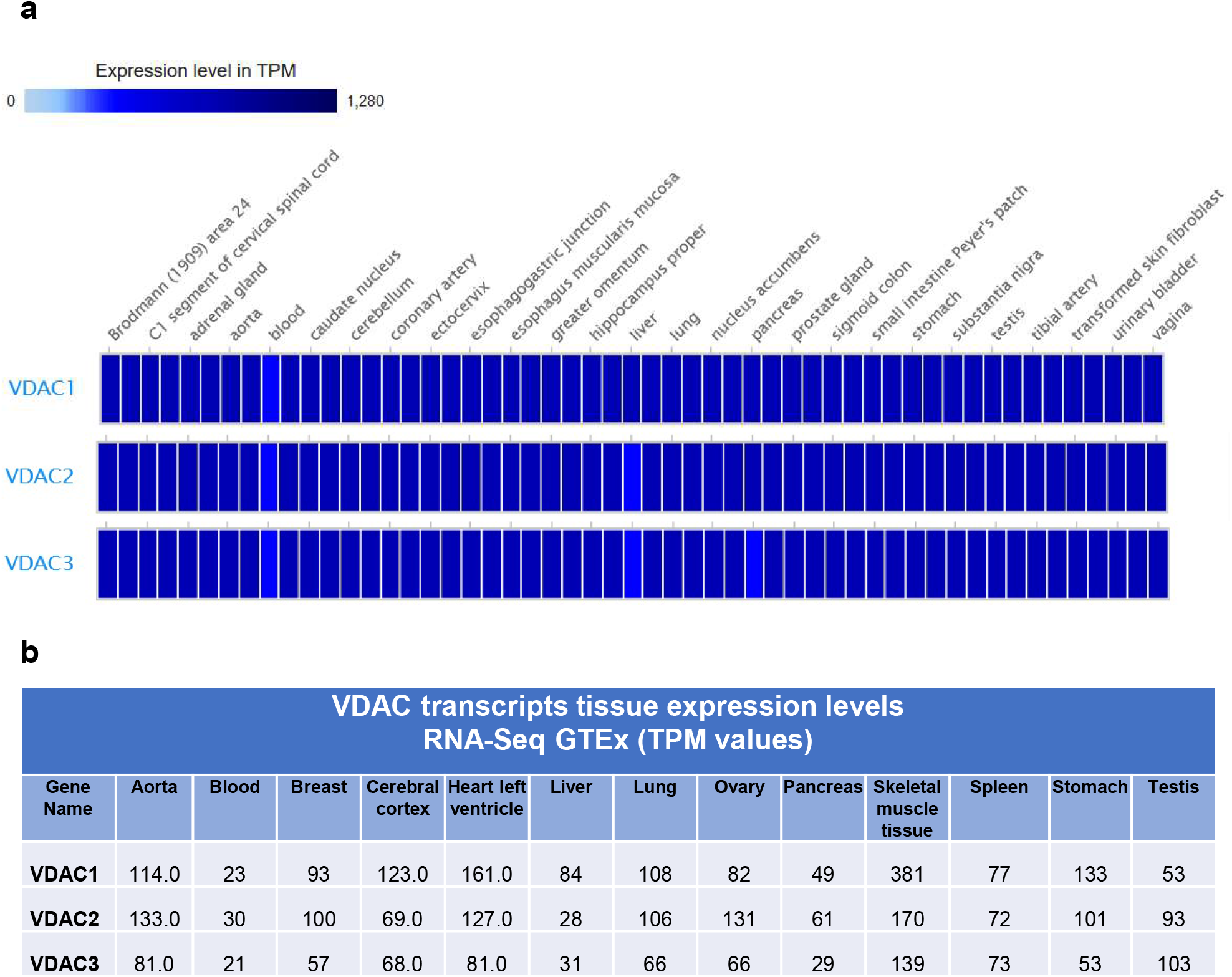
Human VDAC genes expression by Genotype-Tissues-Expression (GTEx) database. RNA-seq data from Genotype-Tissues-Expression (GTEx) project were collected from Gene Expression Atlas repository where all data are manually curated and subject to standardized analysis pipelines. The data are displayed in a heatmap (**a**) with different colours representing a range of TPM mRNA expression level: grey (expression level is below cutoff - 0.5 TPM or FPKM); light blue (expression level is low - between 0.5 to 10 TPM or FPKM); medium blue (expression level is medium - between 11 to 1000 TPM or FPKM); dark blue (expression level is high - more than 1000 TPM or FPKM); white box (no data available). In the panel **b** the specific expression values of each human VDAC isoforms, for representative tissues, are reported.

RNA-seq CAGE VDAC transcripts expression were selected from RIKEN FANTOM 5 project. The enrichment of mRNA obtained by this technique is well known to provide the map of TSS and the activity of genes promoter. In Figure 3 the first relevant result, emerging from this analysis, is the higher expression level of VDAC3 transcripts compared to VDAC1 and VDAC2 in all the human tissues tested. Indeed, VDAC3 mRNA expression falls within a range of TPM (Transcripts Per Kilobase Million) transcripts values higher than that of the other two isoforms. Even if the expression of VDAC1 and VDAC2 transcripts falls in the same range, the latter isoform is the most expressed. In Figure 2 VDACs mRNA levels are reported for some representative tissues. According to the literature and other databases of transcripts expression, VDAC1 and VDAC2 are mainly represented in bone marrow, brain, testis, heart, tongue. However, in FANTOM 5 project dataset the VDAC2 level is doubled compared to VDAC1. Although VDAC3 mRNA expression overcomes VDAC1 and VDAC2, the tissues with a higher level of its expression are confirmed to be heart, testis, muscles. The data emerging from this analysis highlight for the first time the prevalence of VDAC3 gene transcription compared to other isoforms reflecting a higher promoter activity.

**Figure 3.**
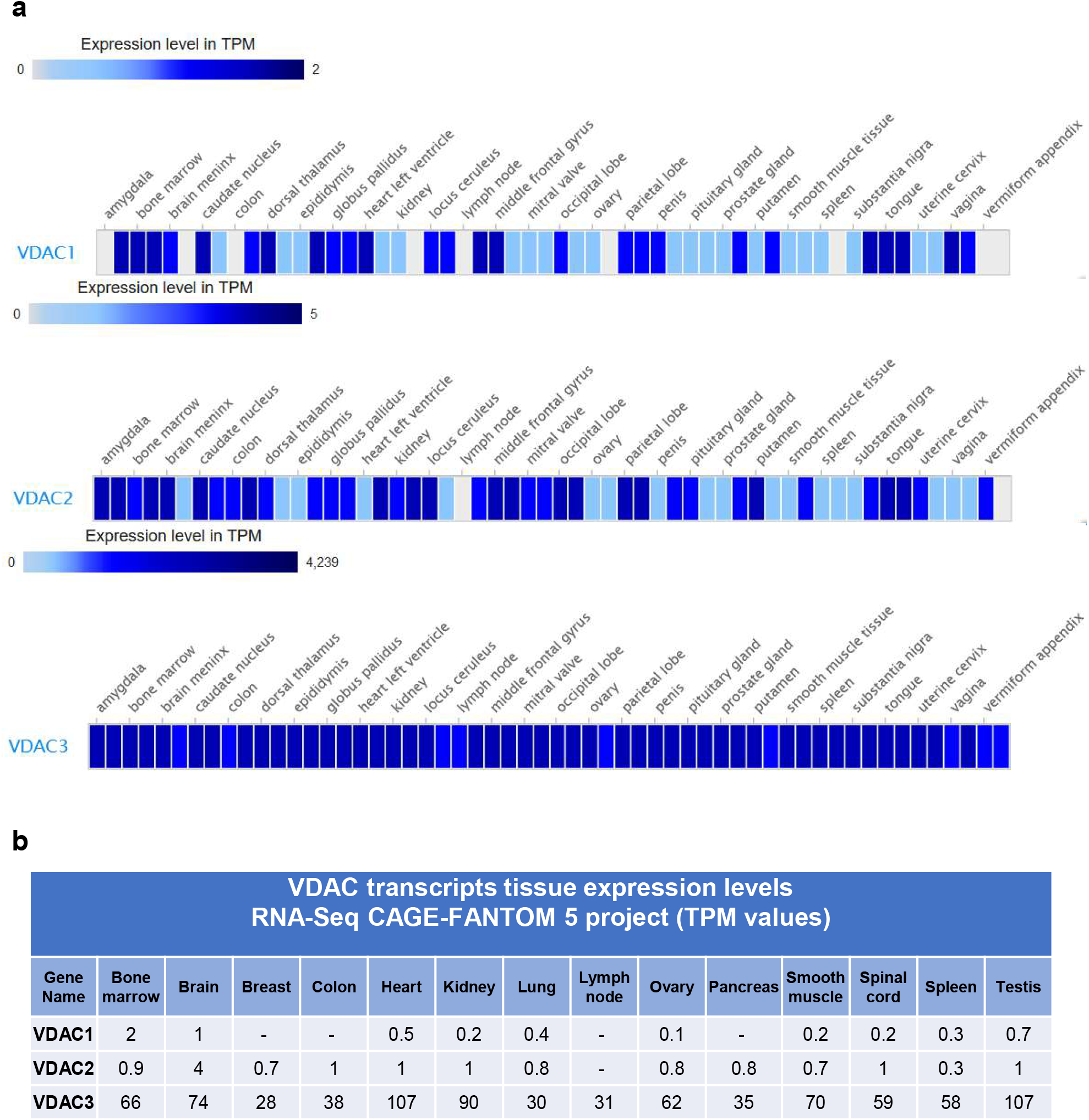
Human VDAC genes expression by RIKEN FANTOM 5 project. RNA-seq CAGE data from RIKEN FANTOM 5 project were collected from Gene Expression Atlas repository where all data are manually curated and subject to standardized analysis pipelines. The data are displayed in a heatmap (**a**) with different colours representing a range of TPM mRNA expression level as described for Figure 2. In the Panel **b** the specific expression values of each human VDAC isoforms, for representative tissues, are reported.

With this analysis, we can also confirm that VDAC isoforms are ubiquitously expressed in tissues even if with different specificity for each isoform.

### VDAC isoforms comparative expression in HeLa cells

VDAC isoforms transcription was analyzed in HeLa cells and a comparison of their expression level was performed. mRNA of VDAC1 gene was established as reference gene and quantification of VDAC2 and VDAC3 mRNA level was reported relative to VDAC1. In Figure 4 a, VDAC2 transcript amount is slightly lower than VDAC1 mRNA showing a value of 0.78, while VDAC3 transcripts are almost half of VDAC1 with a value of 0.39. VDAC3 mRNA is the less expressed transcript among the three isoforms. The data obtained confirm VDAC transcription expression as revealed by GTEx data analysis and previous experimental results obtained by us [34].

**Figure 4.**
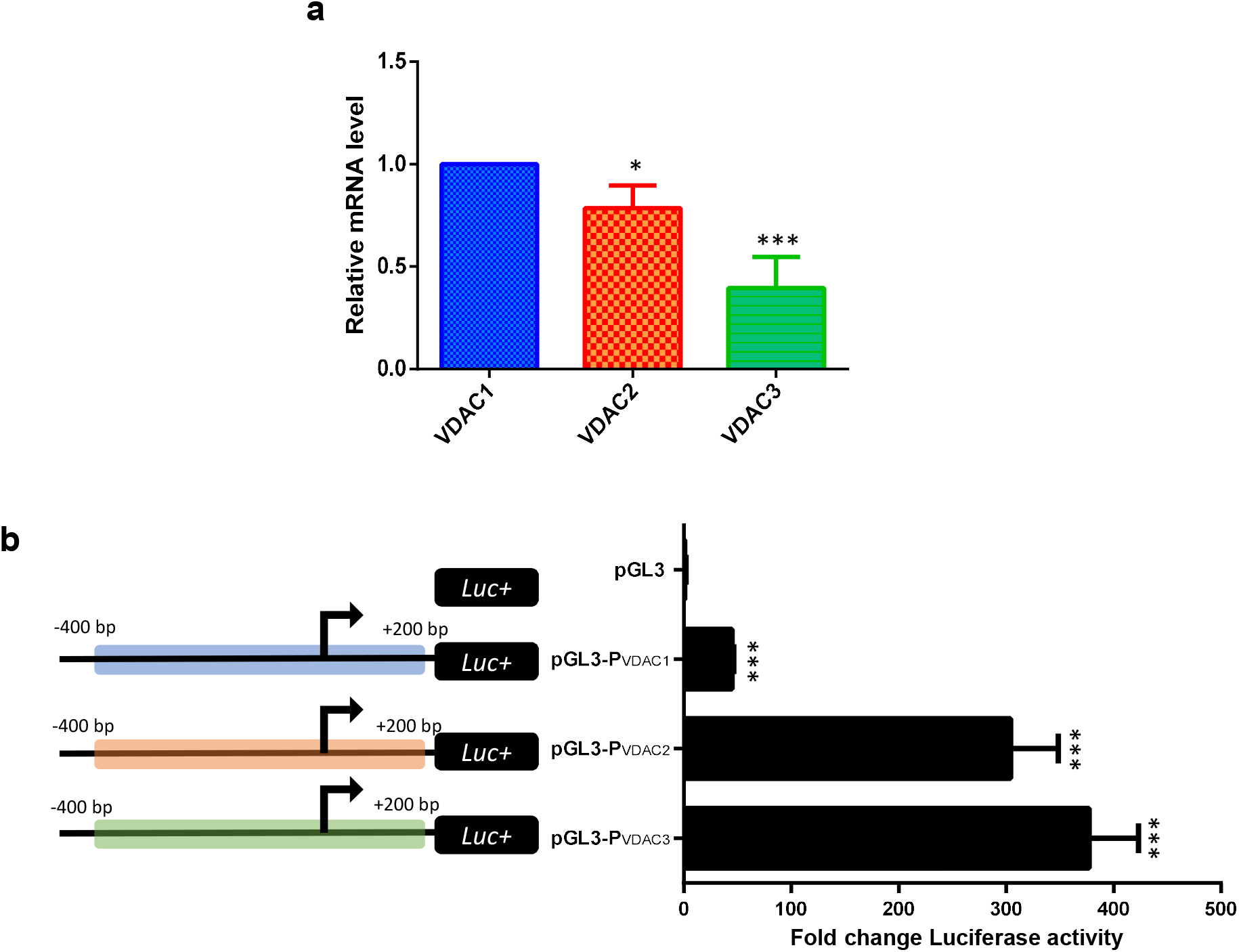
Experimental analysis of human VDAC gene expression and their promoter activity in HeLa cells. **a)** Human VDAC genes expression. VDAC genes expression was detected by Real Time PCR as described in methods. After normalization with the housekeeping gene b-actin, the variation of hVDAC2 and hVDAC3 transcripts was expressed using human VDAC1 as reference. The ∆∆Ct method was applied. **b)** VDAC promoters activity detection. To study the promoters activity, 600 bp sequence encompassing the TSS (from −400 to +200 in the gene sequence) were placed upstream the luciferase gene in pGL3 plasmid. The assay was performed in HeLa cells transfected with P_vdac1_-pGL3, P_vdac2_-pGL3, P_vdac3_-pGL3 constructs after 48h of transfection. Luciferase activity of cell lysates was calculated by referring to empty-pGL3 transfected cells and following normalization with *Renilla* activity. Three independent experiments were performed and results statistically analyzed by one-way ANOVA. A value of P<0,05 was taken as significant.

### VDACs genes transcriptional promoter activity

A 600 bp genomic region encompassing the TSS of VDACs gene was identified as putative promoter named P_*VDAC1*_, P_*VDAC2*_ and P_*VDAC3*_ and utilized for experimental characterization. These sequences were cloned in front of the Luciferase (Luc) reporter gene, to study the activity of the human VDACs genes promoters in HeLa cells. Luciferase activity, driven by the indicated VDACs promoters was compared among the three isoforms as shown in the histogram of Figure 4 b. Surprisingly VDAC1, the most represented isoform holds the less active promoter which drives a transcriptional activation 10 and 8 folds lower than VDAC3 and VDAC2 promoters. These experimental results are in agreement with the predominant transcriptional activity of VDAC3 and VDAC2 emerging from FANTOM 5 project suggesting a mechanism of fine regulation of VDACs expression.

### Characterization of VDAC genes core promoters

Very limited information is available regarding the core promoter organization of VDACs genes. Using different predictive strategies through EPD, YAPP Eukaryotic Core Promoter Predictor and ElemeNT, we built an overview of the most relevant core promoter elements captured with the higher consensus match and functionally recommended scores (p-value ≥ 0,001), which we schematically represented for P_*VDAC1*_ (a), P_*VDAC2*_ (b), and P_*VDAC3*_ (c) (Fig. 5). The promoter region of each VDAC isoform lacks a canonical TATA box, but contains the Initiator element (Inr), downstream promoter element (DPE) and B recognition element (BRE). As typically observed in TATA-less promoters, multiple GC-boxes are required and Inr and DPE are functionally analogous to the TATA box as they cooperate for the binding of TFIID in the transcription [35]. In P_*VDAC2*_ (a) and P_*VDAC3*_ (c) a non-canonical initiation site termed the TCT motif (polypyrimidine initiator) was identified. This polypyrimidine stretch proximal to the 5′ end of these genes is a target for translation regulation and oxidative and metabolic stress, or cancer-induced differential translational regulation by the mTOR pathway [36].

**Figure 5.**
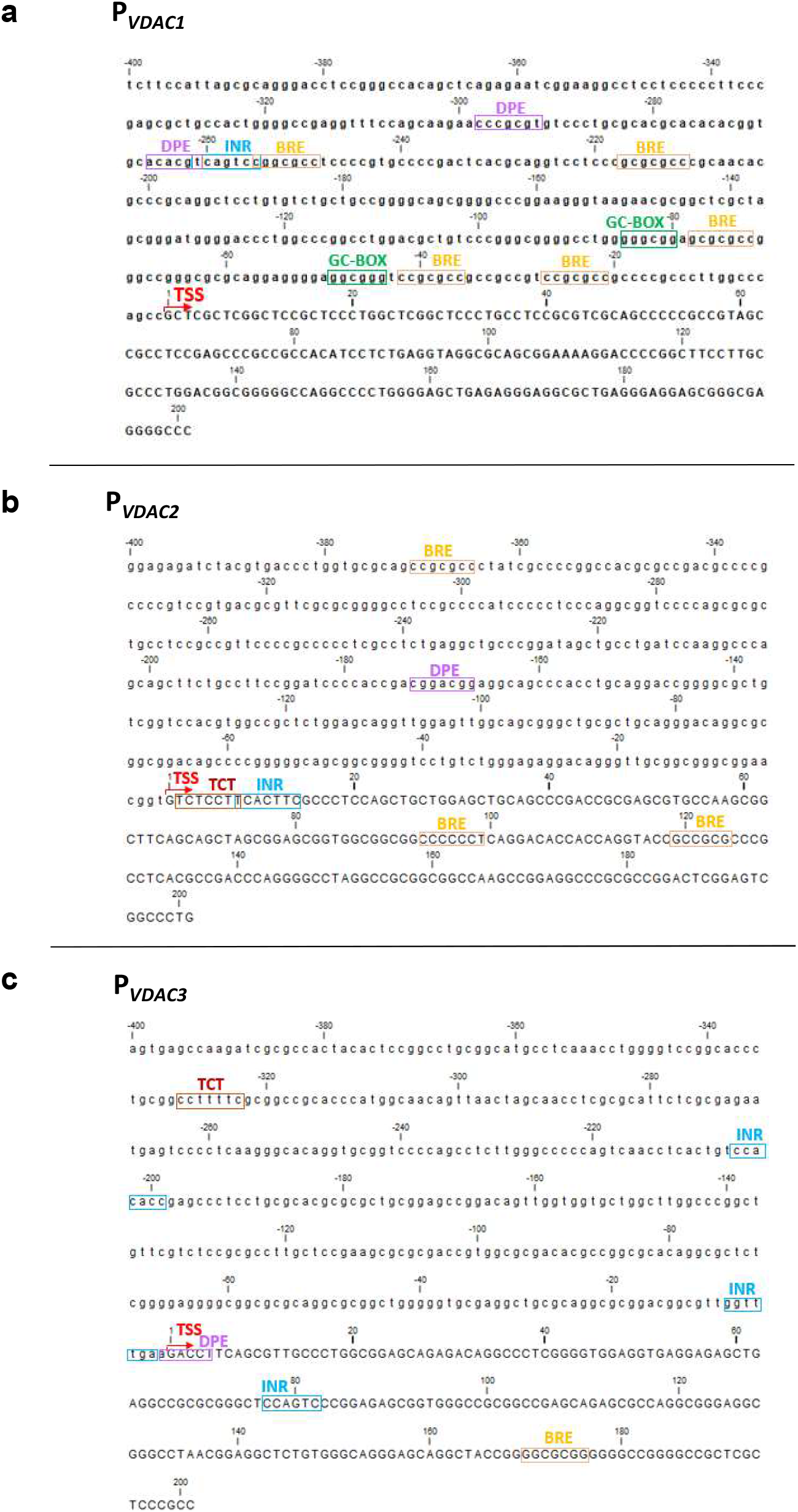
Canonical core promoter elements of human *VDAC* isoforms. The results of core promoter elements identified for *hVDAC1*, *hVD*AC2 and *hVDAC3* genes by predictive tools are the sequence stretches with a high scoring consensus based on position weight matrix (PWM). (**a**) P_*VDAC1*_ (chr5:133,340,230-133,340,830; hg19) encompasses an Inr element (at −261 bp), two GC-boxes (at - 85 bp; −49 bp), five BRE motifs (at −255 bp; −217 bp; −78 bp; −42 bp; −27 bp), and two DPE motifs (at −298 bp; −266). (**b**) P_*VDAC2*_ (chr10:76,970,184-76,970,784; hg19) encompasses a TCT motif, as alternative Inr (at +2 bp), Inr element (at +8 bp), two BRE motifs (at + 93 bp; +119 bp), a DPE motif (at −173 bp). (**c**) P_*VDAC3*_ (chr8:42,248,998-42,249,598; hg19) encompasses three Inr elements (at −205 bp; − 8 bp; + 77 bp), a TCT (at −329 bp), a DPE (at −1 bp) and a BRE (at + 170 bp). TSS site is indicated by a thick red arrow. Nucleotide sequence before TSS is shown in lowercase.

### Characterization of VDACs’ transcriptional regulators

Identifying the upstream regulators of VDAC genes will allow a better understanding of the biological role that each isoform plays in the cell. Thus, VDAC genes were characterized for the TFBSs by scanning the promoter sequences with three different bioinformatics tools (Genomatix, Jaspar, UniBind) and the results were overlapped in order to find the most relevant TF families that regulate VDAC genes expression. We used a search window of −400 to +200 bp around the TSS.

In Figure 6 a, histogram is reported showing every TFBS family found on the VDAC promoters’ sequences that were predicted by MatInspector software (Genomatix v3.10) and experimentally validated by ChIP-Seq data (ENCODE project v3). The ChIP-Seq Peaks of V$E2FF, V$EGRF, V$KLFS, V$NRF1, V$MAZF, V$SP1F, V$ZF02, V$ZF5F are numerically the most overrepresented as highlighted by the bioinformatic prediction, confirming the importance of these factors in the regulatory network of VDAC genes. These binding sites are known to participate in several biological processes, such as cell growth, proliferation and differentiation, development, inflammation and tumorigenesis [37–43]. Among them, V$NRF1 is the master regulator of genes encoding mitochondrial proteins and V$E2FF is a family of factors involved in the control of the cell cycle. The results of Figure 6 a were reorganized in a Venn chart showing the transcription factors binding sites shared by VDACs promoter sequences and those exclusively present in each VDAC gene promoter, as reported by Genomatix analysis (Fig.6 b).

**Figure 6.**
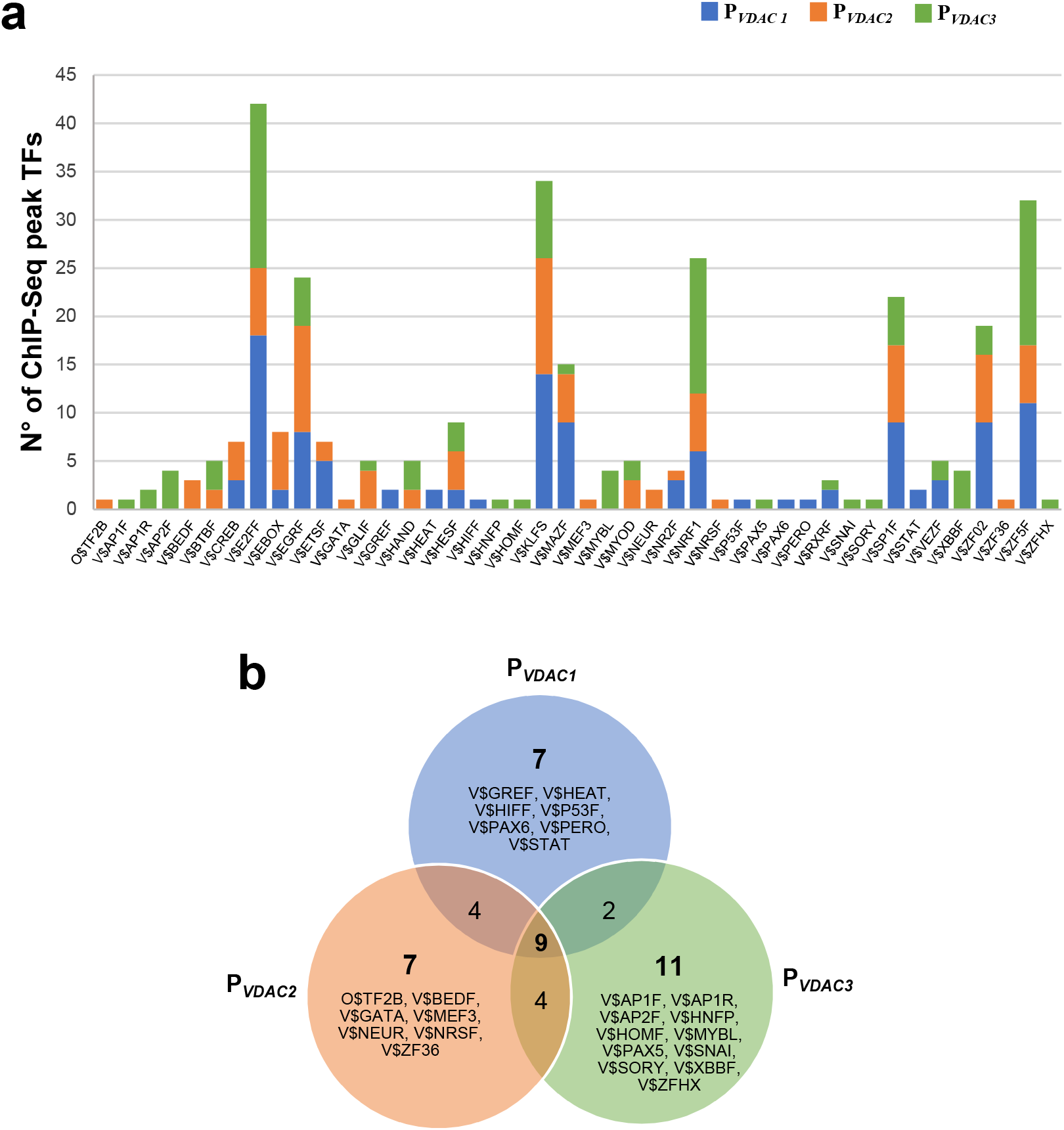
Identification of ChIP-seq peak regions in the human *VDAC* promoters. **a**) The histogram shows the number of TFBS experimentally validated by ChIP-Seq data (ENCODE project v3) among those predicted by the software Genomatix (MatInspector) in P_*VDAC1*_, P_*VDA*C2_ and P_*VDAC3*_ sequences. **b**) Venn diagram showing the number of common and unique predicted binding sites that overlap with a ChIP-Seq region in P_VDAC1_, P_VDAC2_ and P_VDAC3_, based on Genomatix analysis.

#### Analysis of VDACs common TFBSs

The overlap between Genomatix results and data extracted from JASPAR and UniBind databases highlights the occurrence of four families of shared TFBSs in promoter regions of VDAC genes. A comprehensive location of both sets of TFBSs was also extracted from ENCODE ChIP-seq peaks (Table 1). The detection of common TFBS clusters indicates that these different classes of TFs participate in many similar activities prevalently involved in cell proliferation and differentiation, apoptosis and metabolism regulation. Therefore, it is possible to divide different classes of TFs involved in the control of VDAC genes in three functional categories: the first one is represented by V$E2FF (E2F-myc activator/cell cycle) transcription factors, affecting various processes of cell cycle regulation [44]. The second group includes members of V$EBOX (E-box binding factors) and V$KLFS (Krueppel like transcription factors) families, which are essential transcription factors that regulate a large number of cellular processes, such as metabolism, cell proliferation, differentiation, apoptosis, and cell transformation [45, 46]. The third group comprises V$NRF1 (Nuclear respiratory factor 1) family, closely connected with mitochondrial biogenesis, DNA damage signalling, and tumour metabolism [47–49].

**Table 1.** VDACs common TFBSs

| Matrix Family | Description | RATE OF FREQUENCY IN DB |  |  |  |  |  |  |  |  |
| --- | --- | --- | --- | --- | --- | --- | --- | --- | --- | --- |
|  |  | P<sub>VDAC1</sub> |  |  | P<sub>VDAC2</sub> |  |  | P<sub>VDAC3</sub> |  |  |
|  |  | Genomatix | Jaspar | Unibind | Genomatix | Jaspar | Unibind | Genomatix | Jaspar | Unibind |
| V\$E2FF | E2F-myc activator/cell cycle regulator | ✓ | ✓ | ✓ | ✓ | ✓ | ✓ | ✓ | ✓ | - |
| V\$EBOX | E-box binding factors | ✓ | ✓ | ✓ | ✓ | - | - | ✓ | ✓ | ✓ |
| V\$KLFS | Krueppel like transcription factors | ✓ | ✓ | ✓ | ✓ | ✓ | ✓ | ✓ | ✓ | ✓ |
| V\$NRF1 | Nuclear respiratory factor 1 | ✓ | ✓ | ✓ | ✓ | ✓ | ✓ | ✓ | ✓ | ✓ |

#### Analysis of VDACs unique TFBSs

The data extracted from Genomatix, Jaspar, and UniBind, were collected to obtain information about families of TFBS exclusively found in each promoter sequence: they thus define the unique TFs controlling each single VDAC isoform. We found in the P_*VDAC1*_ sequence (Table 2) four unique TFBS families: V$AHRR (AHR-arnt heterodimers and AHR-related factors) which is required, together with HIF-1α factor, for the cell response to hypoxia [50]; V$ETSF (Human and murine ETS1 factors) that includes NRF2, a regulator of mitochondrial biogenesis and redox homeostasis [51]; V$HEAT (heat shock factors) a family of proteins crucial for cell stress response [52]; V$PBXC (PBX-MEIS complexes), also known as pre-B cell leukemia family, that includes regulators of cell development, survival, invasion and proliferation [53].

**Table 2.** VDAC1 unique TFBSs

| Matrix Family | Description | RATE OF FREQUENCY IN DB |  |  |
| --- | --- | --- | --- | --- |
| | | $P_{VDAC1}$ | | |
|  |  | *Genomatix* | *Jaspar* | *UniBind* |
| V\$AHRR | AHR-arnt heterodimers and AHR-related factors | - | - | ✓ |
| V\$ETSF | Human and murine ETS1 factors | ✓ | - | - |
| V\$HEAT | Heat shock factors | ✓ | - | - |
| V\$PBXC | PBX - MEIS complexes | - | - | ✓ |

As concerning P_*VDAC2*_ sequence (Table 3), we identified several binding sites recognized by regulators know to be associated with the nervous system development and general core promoter elements. Among them: V$BEDF (BED subclass of zinc-finger proteins) includes ZBED which controls cell growth and differentiation in cone photoreceptors and Müller cells of human retina [54]; V$BRAC (Brachyury gene, mesoderm developmental factor), is involved in the commitment of T helper (Th) cells [55]; V$CLOX (CLOX and CLOX homology (CDP) factors), a crucial regulator of the neuronal differentiation in the brain [56]; V$MEF3 (MEF3 binding sites) a family that includes regulators of skeletal muscle development [57]; V$NEUR (NeuroD, Beta2, HLH domain), comprising the basic helix-loop-helix factors Ascl1 and OLIG2 involved in neural development and differentiation [58]; TF2B (RNA polymerase II transcription factor II B) a core promoter element [59]; V$ZFXY (Zfx and Zfy - transcription factors), a family of transcription factors implicated in mammalian sex de Termination [60].

**Table 3.**
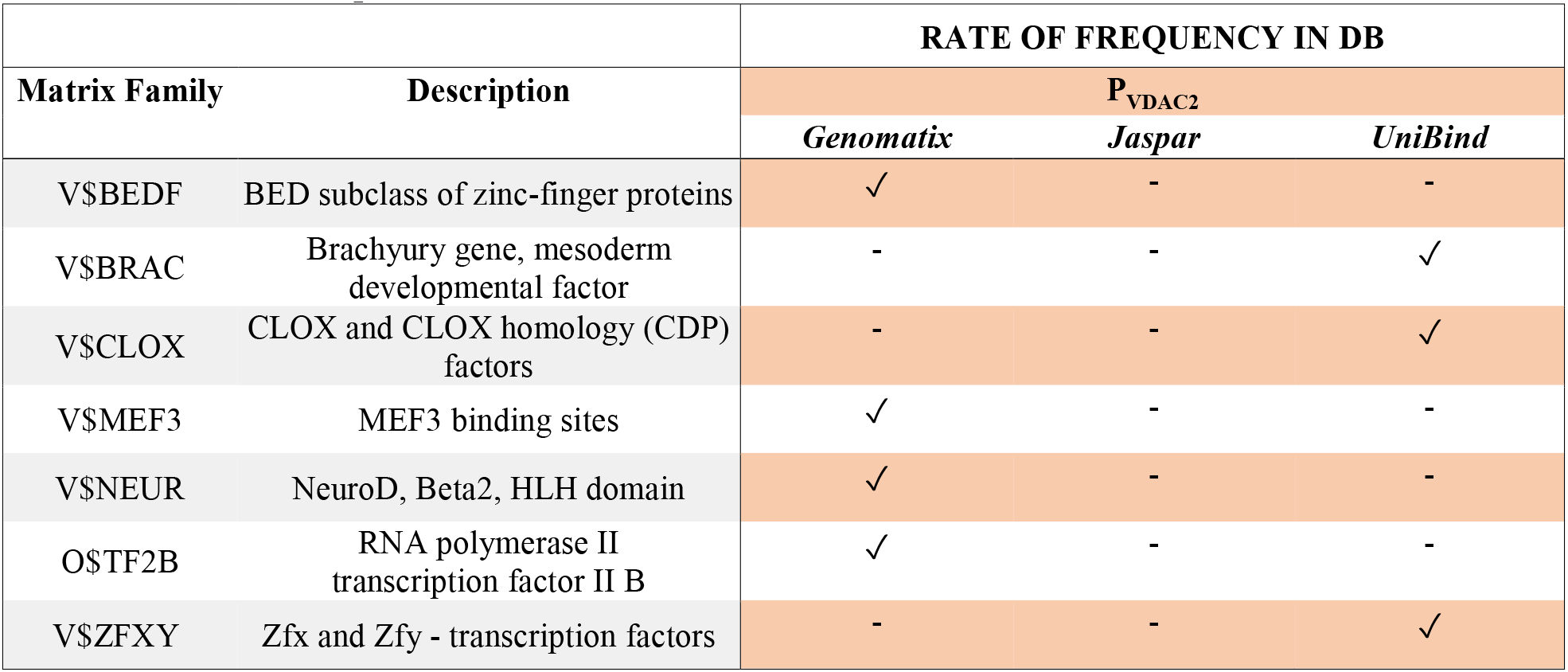
VDAC2 unique TFBSs

The results of P_*VDAC3*_ analysis (Table 4) showed a distribution of binding sites for TFs involved in the control of various cellular processes including cell differentiation, proliferation, apoptosis, and gametogenesis: V$BCL6 (BED subclass of zinc-finger proteins), a critical regulator of B cell differentiation [61]; V$CDXF (Vertebrate caudal related homeodomain protein) involved in development and maintenance of trophectoderm [62]; V$FOX (Forkhead (FKH)/ Forkhead box (Fox)), including important regulators of development, organogenesis, metabolism and cell homeostasis [63]; V$SOHLH (Spermatogenesis and oogenesis basic helix-loop-helix) transcription regulators of male and female germline differentiation [64]; V$HMG (High-Mobility Group family), including factors that regulate neuronal differentiation and also play important roles in tumorigenesis [65]; V$HOMF (Homeodomain transcription factors) involved in central nervous development [66]; V$IRFF (Interferon regulatory factors) required for differentiation of hematopoietic cells [67]; V$LBXF (Ladybird homeobox (lbx) gene family) that plays a critical role in embryonic neurogenesis and myogenesis and in muscle mass determination [68]; V$MYBL (cellular and viral myb-like transcriptional regulators) that controls cell cycle progression, survival and differentiation [69]; V$SMAD (Vertebrate SMAD family of transcription factors) that includes factors responsible of several cellular processes, including proliferation, differentiation, apoptosis, migration, as well as cancer initiation and progression [70]; V$XBBF (X-box binding factors) family involved in the control of development and maintenance of the endoplasmic reticulum (ER) in multiple secretory cell lineages [71].

**Table 4.**
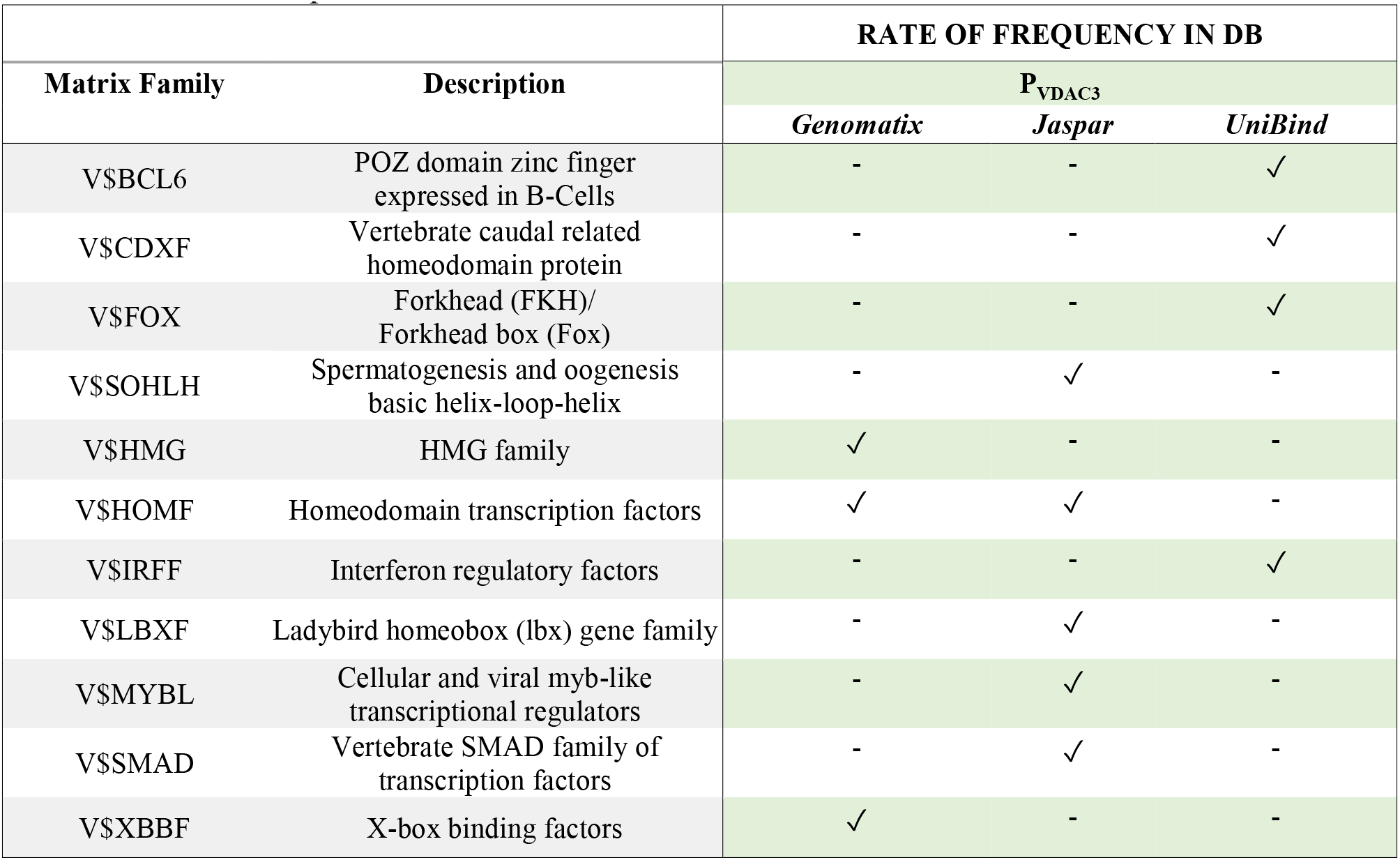
VDAC3 unique TFBSs

In Figure 7 (a-c) a magnification of VDACs promoters analyzed from UCSC Genome Browser is overlapped with experimental data proving the transcriptional activity of this genomic region. Based on Genomatix results on distinct and shared TFBSs at promoter regions of P_*VDAC1*_, P_*VDAC2*_, and P_*VDAC3*_, a comprehensive location of both sets of TFBSs was extracted from ENCODE ChIP-seq peaks. These findings are also supported by TFBSs enrichment analyses from JASPAR and UniBind database. In the graphical view, the most interesting peaks of TFBS found, are located in overlapping positions of the promoter for different cell lines. Moreover, these validated TFBSs fall in the genomic region corresponding to VDACs promoter studied in this work.

**Figure 7.**
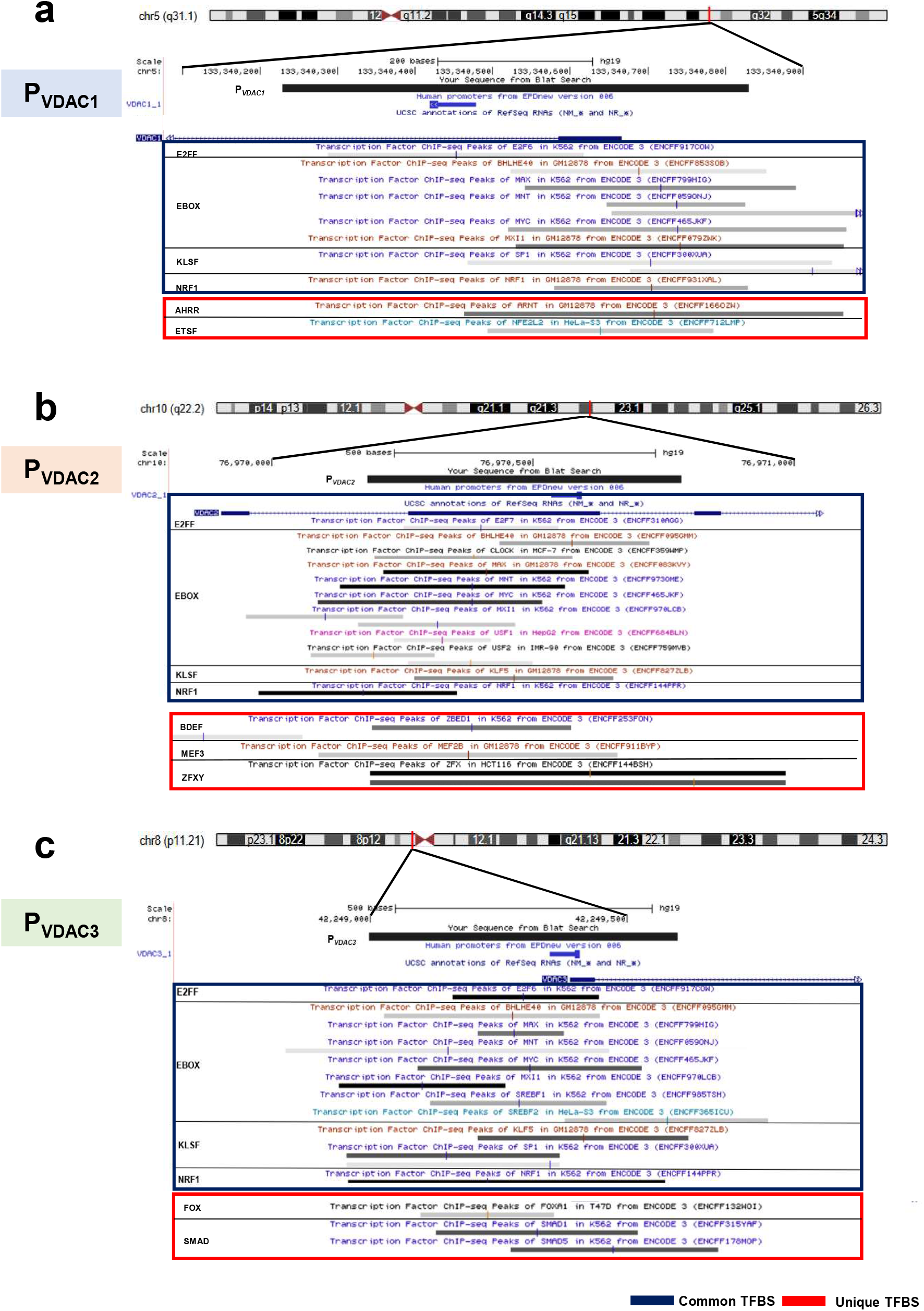
Identification of common and unique transcription factor binding site clusters of *VDACs* promoter sequences. Distribution of a common set (enclosed in a blue box) and specific sets (enclosed in a red box) of Transcription Factors binding sites (TFBS) in VDAC isoforms promoters as reported in different cell lines by ChIP-Seq analysis in ENCODE Project (shown as colour vertical bars in the gray segments). **a**) *hVDAC1*, **b**) *hVDAC2*, and **c**) *hVDAC3*.

The determination of common TFBSs appear to corroborate shared biological properties, as well as a high degree of functional conservation and cooperation among the three isoforms, while the mapping of unique TFBSs robustly supported by different databases suggest a different biological role.

## Discussion

To understand the specialized biological role of VDACs isoforms, simultaneously expressed in cells, we performed a characterization of VDACs transcripts expression and promoters’ structure and function. To have a general but reliable picture of VDAC genes structure, expression, and regulation, we undertook a study of VDACs isoforms in the main available public resource reporting high throughput data of international collaborative projects.

### Structure of VDAC genes, transcripts and promoters

First of all, a general view of VDAC genes, transcripts variants, and promoter regions feature by in silico analysis through UCSC genome browser is shown. For each VDAC genes, several different transcripts splice variants were identified varying not in the coding region but mainly in their 5’-UTR and 3’-UTR length. Other variants are processed transcripts, other present retained intron and for VDAC3 two are involved in nonsense-mediated decay mechanism. The variability on the UTR sequences let to hypothesize differentiated mechanism of transcript regulation and expression context for each variant. The 3’-UTR sequence may be a target of translation regulation by miRNA or interference. Many publications indeed reported the identification of miRNA molecules targeting all three VDACs transcripts but in particular VDAC1 [72]. The 5’-UTR region variability might be associated with alternative promoter usage and activation in a different expression context. However, no information is available on the human VDACs promoter and/or other regulative regions to explain transcripts expression.

### VDAC expression in Gene Expression Atlas repository

For this reason, we wanted to focus our study on the characterization of the main promoter driving each VDACs genes transcripts expression. First of all, we selected from Gene Expression Atlas repository, the data derived from the RNA-seq CAGE RIKEN FANTOM 5’ project and the RNA-seq GTEx projects.

Generally, although the expression profile of VDACs transcripts presents a differentiated level in different tissues or cells types, all three isoforms are ubiquitously expressed [2]. The level of VDAC1 and VDAC2 transcripts is comparable, while VDAC3 is always less expressed [34]. Surprisingly, analyzing the data set of FANTOM 5 project we found that VDAC3 transcripts expression overcome VDAC2 but in particular VDAC1 which results to be scarcely represented in all tissues compared to the other VDACs. The special version of RNA-seq methodology based on cap analysis of gene expression adopted by the FANTOM5 consortium allowed to identify active TSS located on the 5’-end of transcribed mRNA which are not necessarily associated to a full length and/or protein coding transcript. Based on this evidence, we selected the main promoter region found in Eukaryotic Promoter Database EPD associated to the main protein-coding transcript and we confirmed by luciferase reporter assay the highest transcriptional activity of VDAC3 promoter and the VDAC1 which is on the contrary the less active. These interesting results suggest a mechanism of more complex and coordinated regulation of VDACs transcripts expression in order to enhance the most represented and functional VDAC1 isoform and/or repress VDAC3. However, the higher promoter activity of VDAC3 might be also interpreted as a potentiality maintained by the cells to promptly respond to a particular and still unknown stimulation through VDAC3 increased expression in specific conditions.

### VDAC genes core promoters

With the aim to explore the mechanism of VDAC transcription regulation, we started a systematic analysis of human VDAC genes promoters to highlight their structural and functional features. VDACs genes core promoter organization is similar to most of TATA-less human core promoters of ubiquitously expressed genes. Abundant GC regions, alternative binding sites Inr, DPE and BRE for the basal transcription factors take over the function of TATA box sequence. Moreover, VDACs genes, as most of human protein-coding genes lacking TATA-box, own a long 5’-UTR region suggesting the presence of alternative TSS employed for expression of distinct products in different contexts or tissues. As reported in the VDACs genes structure organization, the occurrence of several transcripts, could explain their expression associated with different conditions.

### Transcription factors binding sites common to any VDAC gene

We also characterized the main transcription factors regulating the activity of VDAC promoter regions, looking for the transcription factors binding sites (TFBSs). The information we gained by bioinformatic analysis, suggested the central role of VDAC protein expression in regulating mitochondrial function in fundamental cell processes. We recognized TFBS shared by the three VDACs promoters, as well as single promoters’ unique sites. Among the common ones, the majority of identified TFs classes belong to the E2FF, NRF1, SP1, KLFS, EBOX families which participate in many similar activities but are prevalently involved in cell proliferation and differentiation, apoptosis and metabolism regulation [37–43]. TFBS for E2FF and NRF1 transcription factor family members are also numerically the most represented on all the three VDACs promoters. VDACs promoters are mainly characterized by a large number of recognition sites for E2FF and NRF1 transcription factors, which were found associated by chromatin immunoprecipitations with microarrays (ChIP-on-chip) to a significant subset of genes implicated in mitochondrial biogenesis and metabolism, other than mitosis, cytokinesis, cell cycle control, grow, proliferation [73]. Many identified TFBS are located in the proximal core promoter acting as co-regulators for general transcription factors activation and chromatin regulation. In particular, some of them, SP1, KLFS, EBOX, NRF1 rich in GC content have an important role in epigenetic control of promoter suggesting a more complex regulation of this genes [74].

### Transcription factors binding sites specific to each VDAC gene

Search for unique transcription factor binding sites on the promoters of VDAC isoforms, allowed us to hypothesize their involvement in specialized biological functions. However, the transcription factors specific for each VDAC genes are always correlated to essential functions ensuring cell survival and functionality. In general, these processes require a noteworthy energy cost: metabolism maintenance, development, organogenesis, dysfunction of mitochondria in pathology, are some examples.

The families of transcription regulators identified as unique in VDAC1 promoter suggests that this isoform was probably selected by evolutionary process to have the prevalent role of channel protein in the mitochondrial outer membrane in physiological context and in particular when altered conditions force the cells to restore the mitochondria energetic balance [75]. These observations are corroborated by several experimental evidence showing the involvement of VDAC1 in regulating many cellular and mitochondrial events in pathology or stress conditions through the interaction with specific protein [76].

VDAC2 was indicated as the isoform carrying out channel function and governing apoptosis and autophagy in various contexts [5]. Analysis of VDAC2 promoter highlighted the presence of different factors specially involved in development of specialized tissues and organogenesis process as unique among VDACs promoters. Most of these factors are related to nervous system genesis and development.

VDAC3 is controlled by the most active promoter: it is particularly rich of GC repetitions, suggesting an epigenetic control mechanism able to reduce the expression of transcripts. Factors binding sites found in VDAC3 promoter belong to various families but those involved in development of germinal tissues, organogenesis and sex determination are the most abundant. Also in this case, the experimental evidence reported in the literature confirms the crucial role of VDAC3 in fertility [13].

## Conclusion

In conclusion we proposed a general overview of the structural and functional organization of VDACs isoforms promoters cross-referencing public available data source, bioinformatics prediction and experimental data. From this analysis, it emerges the essential function of the family of VDAC proteins in the regulation of mitochondrial energetic metabolism in physiological and pathological cell life. Moreover, we shed some new light on the molecular mechanisms that explain the differences among three VDAC isoforms. It is becoming increasingly clear that the most known specialized functions of each VDAC isoforms are connected with the organization of the “button room” that decides the transcriptional activity of their genes and were produced by evolution.

## Materials and Methods

### Bioinformatic analysis of promoter region

Human promoter retrieval for *hVDAC1* (NM_003374), *hVDAC2* (NM_001324088) and *hVDAC3* (NM_005662) genes was carried out by high-quality promoter resource EPDnew version 006 (https://epd.epfl.ch). For study purposes, P_*VDAC1*_ (chr5:133,340,230-133,340,830; hg19) P_*VDAC2*_ (chr10:76,970,184-76,970,784; hg19) and P_*VDAC3*_ (chr8:42,248,998-42,249,598; hg19) are the promoter sequences extended from −400 bp to +200 bp relative to annotated Transcription Start Site (TSS) of basal EPD promoter sequences (VDAC1_1; VDAC2_1; VDAC3_1). Tools used to scan for canonical core promoter elements and synergistic combinations were EPD Promoter Elements Page, YAPP Eukaryotic Core Promoter Predictor (http://www.bioinformatics.org/yapp/cgi-bin/yapp_intro.cgi), and ElemeNT (http://lifefaculty.biu.ac.il/gershon-tamar/index.php/resources). Analysis of Transcription Factor Binding Site (TFBS) clusters was done by Genomatix software suite (Genomatix v3.10). *MatInspector* application was used to identify potential binding sites for transcription factors (TFBSs) in input sequence using the Matrix Family Library version 11.0 for core promoter elements in vertebrates with a fixed matrix similarity threshold of 0.85 [77].

TFBSs enrichment analyses was also performed using JASPAR database, a carefully selected DB of position-specific scoring matrices derived from experimentally validated TFBS [78], and UniBind database encompassing predicted direct TF–DNA interactions derived from PWMs covering >2% of the human genome [79]. In view of implementing the TFBS clusters analysis, all information relevant to the genomic structure and DNA regulatory elements related to three VDAC isoforms were investigated using UCSC Genome Browser (https://genome.ucsc.edu). DNA regulation tracks of UCSC Genome Browser and some information of promoter sequence and TFBS predictions are currently available on GRCh37/hg19.

### Gene expression data retrieval

Gene expression results were collected from Gene Expression Atlas of EMBL-EBI open science resource (https://www.ebi.ac.uk/gxa/home), a database of RNA-seq, microarray and, proteomics data manually curated and analyzed through standardized analysis pipelines. The Baseline Atlas database containing the RNA-seq experiments regarding expression of gene in tissues under physiological conditions was consulted and data from Genotype-Tissue-Expression (GTEx) [32] and Functional ANnotation of the Mammalian Genome 5 (FANTOM) project [33] were selected for VDAC isoforms expression analysis. The data are displayed in a heatmap with different colour representing a range of TPM mRNA expression level. Transcripts expression level of the three VDACs gene were selected for the most common tissues and represented by a histogram and reported in a table.

### Quantitative Real-time PCR

0.6 ×10^6^ of HeLa cells were plated in 25 cm^2^ flasks. After 48h of incubation total RNA was extracted using “ReliaPrep RNA cell mini-prep system” (Promega) according to manufacturer’s instructions. RNA concentration and purity were measured by a spectrophotometer and 2 ug were used to synthesize cDNA by QuantiTect Reverse Transcription kit (Qiagen). Real-time amplification was performed in a Mastercycler EP Realplex (Eppendorf) in 96-well plates. The reaction mixture contained 1.5 ul cDNA, 0.2 uM gene specific primers pairs (hVDAC1, hVDAC2, hVDAC3, ß-actin) and 12.5 ul of master mix (QuantiFast SYBR Green PCR kit, Qiagen). Three independent experiments of quantitative real-time were performed in triplicate for each sample. Analysis of relative expression level was performed by the ∆∆C_t_ method using the housekeeping ß-actin gene as internal calibrator and VDAC1 gene as reference.

### Plasmid constructs

A putative promoter region of 600bp encompassing the TSS of the human VDACs genes was selected from GenBank and cloned into pGL3 basic vector (Promega) for transcriptional activity study. The construct P_*VDAC1*_ contain the sequence (chr5:133,340,230-133,340,830; hg19) derived from VDAC1 NM_003374, P_*VDAC2*_ contain the genomic trait (chr10:76,970,184-76,970,784; hg19) from VDAC2 NM_001324088 and P_*VDAC3*_ the sequence (chr8:42,248,998-42,249,598; hg19) of VDAC3 NM_005662.

### Promoter reporter assay

HeLa cells were plated at density of 0.3 × 10^6^ cells/well in a 6 wells plate. After 24h, cells were transfected with 800 ng of pGL3 constructs and 20 ng of pRL-TK renilla reporter vector by Transfast transfection reagent according to manufacturer’s protocol (Promega) and after 48h cells were lysed. Luciferase activity of cell lysate transfected with pGL3 promoter constructs was detected with the Dual Luciferase Assay (Promega) according to the manufacturer’s instructions. Activity of firefly luciferase relative to renilla luciferase was expressed in relative luminescence units (RLU). Variation of luminescence units of treated samples relative to control, were indicated as fold increase (FI).

## Abbreviations

AHRR: AHR-arnt heterodimers and AHR-related factors
BCL6: BED subclass of zinc-finger proteins
BEDF: BED subclass of zinc-finger proteins
BRAC: Brachyury gene, mesoderm developmental factor
BRE: B recognition element
CDXF: Vertebrate caudal related homeodomain protein
ChIP-on-chip: CHromatin ImmunoPrecipitations with microarrays
CLOX: CLOX and CLOX homology (CDP) factors
DPE: Downstream promoter element
E2FF: E2F-myc activator/cell cycle
EBOX: E-box binding factors
ER: Endoplasmic Reticulum
ETSF: Human and murine ETS1 factors
FANTOM: Functional ANnotation of the Mammalian Genome
FOX: Forkhead (FKH)/ Forkhead box (Fox)
GTEX: Genotype-Tissue Expression
HEAT: heat shock factors
HMG: High-Mobility Group family
HOMF: Homeodomain transcription factors
Inr: Initiator Element
IRFF: Interferon regulatory factors
KLFS: Krueppel like transcription factors
LBXF: Ladybird homeobox (lbx) gene family
MEF3: MEF3 binding sites
MYBL: cellular and viral myb-like transcriptional regulators
NEUR: NeuroD, Beta2, HLH domain
NRF1: Nuclear respiratory factor 1
PBXC: PBX-MEIS complexes
SMAD: Vertebrate SMAD family of transcription factors
SOHLH: Spermatogenesis and oogenesis basic helix-loop-helix
TF2B: RNA polymerase II transcription factor II B
TFBSs: Transcription Factor Binding Sites
TSS: Transcription Start Site
VDAC: Voltage-Dependent Anion selective Channel
XBBF: X-box binding factors
ZFXY: Zfx and Zfy - transcription factors

## Notes

### Competing Interest Statement

The authors have declared no competing interest.

## References

1. Shoshan-Barmatz V, De Pinto V, Zweckstetter M, Raviv Z, Keinan N, Arbel N. VDAC, a multi-functional mitochondrial protein regulating cell life and death. Molecular aspects of medicine. 2010 Jun;31(3):227–85. PubMed PMID: 20346371.

2. Messina A, Reina S, Guarino F, De Pinto V. VDAC isoforms in mammals. Biochimica et biophysica acta. 2012 Jun;1818(6):1466–76. PubMed PMID: 22020053.

3. Colombini M. VDAC: the channel at the interface between mitochondria and the cytosol. Molecular and cellular biochemistry. 2004 Jan-Feb;256-257(1-2):107–15. PubMed PMID: 14977174.

4. Benz R. Permeation of Hydrophilic Solutes through Mitochondrial Outer Membranes - Review on Mitochondrial Porins. Bba-Rev Biomembranes. 1994 Jun 29;1197(2):167–96. PubMed PMID: WOS:A1994NX15200004. English.

5. Naghdi S, Hajnoczky G. VDAC2-specific cellular functions and the underlying structure. Biochimica et biophysica acta. 2016 Oct;1863(10):2503–14. PubMed PMID: 27116927. Pubmed Central PMCID: 5092071.

6. Checchetto V, Reina S, Magri A, Szabo I, De Pinto V. Recombinant human voltage dependent anion selective channel isoform 3 (hVDAC3) forms pores with a very small conductance. Cellular physiology and biochemistry : international journal of experimental cellular physiology, biochemistry, and pharmacology. 2014;34(3):842–53. PubMed PMID: 25171321.

7. Queralt-Martin M, Bergdoll L, Teijido O, Munshi N, Jacobs D, Kuszak AJ, et al. A lower affinity to cytosolic proteins reveals VDAC3 isoform-specific role in mitochondrial biology. The Journal of general physiology. 2020 Feb 3;152(2). PubMed PMID: 31935282. Pubmed Central PMCID: 7062508.

8. Sampson MJ, Lovell RS, Davison DB, Craigen WJ. A novel mouse mitochondrial voltage-dependent anion channel gene localizes to chromosome 8. Genomics. 1996 Aug 15;36(1):192–6. PubMed PMID: 8812436.

9. Neumann D, Buckers J, Kastrup L, Hell SW, Jakobs S. Two-color STED microscopy reveals different degrees of colocalization between hexokinase-I and the three human VDAC isoforms. PMC biophysics. 2010 Mar 5;3(1):4. PubMed PMID: 20205711. Pubmed Central PMCID: 2838807.

10. Reina S, Palermo V, Guarnera A, Guarino F, Messina A, Mazzoni C, et al. Swapping of the N-terminus of VDAC1 with VDAC3 restores full activity of the channel and confers anti-aging features to the cell. FEBS letters. 2010 Jul 2;584(13):2837–44. PubMed PMID: 20434446.

11. Anflous K, Armstrong DD, Craigen WJ. Altered mitochondrial sensitivity for ADP and maintenance of creatine-stimulated respiration in oxidative striated muscles from VDAC1-deficient mice. The Journal of biological chemistry. 2001 Jan 19;276(3):1954–60. PubMed PMID: 11044447.

12. Cheng EH, Sheiko TV, Fisher JK, Craigen WJ, Korsmeyer SJ. VDAC2 inhibits BAK activation and mitochondrial apoptosis. Science. 2003 Jul 25;301(5632):513–7. PubMed PMID: 12881569.

13. Sampson MJ, Decker WK, Beaudet AL, Ruitenbeek W, Armstrong D, Hicks MJ, et al. Immotile sperm and infertility in mice lacking mitochondrial voltage-dependent anion channel type 3. The Journal of biological chemistry. 2001 Oct 19;276(42):39206–12. PubMed PMID: 11507092.

14. Menzel VA, Cassara MC, Benz R, de Pinto V, Messina A, Cunsolo V, et al. Molecular and functional characterization of VDAC2 purified from mammal spermatozoa. Bioscience reports. 2009 Jul 22;29(6):351–62. PubMed PMID: 18976238.

15. Manczak M, Sheiko T, Craigen WJ, Reddy PH. Reduced VDAC1 protects against Alzheimer’s disease, mitochondria, and synaptic deficiencies. Journal of Alzheimer’s disease : JAD. 2013;37(4):679–90. PubMed PMID: 23948905. Pubmed Central PMCID: 3925364.

16. Nowak G, Megyesi J, Craigen WJ. Deletion of VDAC1 Hinders Recovery of Mitochondrial and Renal Functions After Acute Kidney Injury. Biomolecules. 2020 Apr 10;10(4). PubMed PMID: 32290153. Pubmed Central PMCID: 7226369.

17. Huizing M, Ruitenbeek W, Thinnes FP, DePinto V, Wendel U, Trijbels FJ, et al. Deficiency of the voltage-dependent anion channel: a novel cause of mitochondriopathy. Pediatric research. 1996 May;39(5):760–5. PubMed PMID: 8726225.

18. Huizing MR, W.; Thinnes, F.; De Pinto, V. Deficiency of the Voltage-Dependent Anion Channel (VDAC): a novel cause of mitochondrial myopathies. The Lancet. 1994:344:762.

19. Baines CP, Kaiser RA, Sheiko T, Craigen WJ, Molkentin JD. Voltage-dependent anion channels are dispensable for mitochondrial-dependent cell death. Nature cell biology. 2007 May;9(5):550–5. PubMed PMID: 17417626. Pubmed Central PMCID: 2680246.

20. Reina S, Pittala MGG, Guarino F, Messina A, De Pinto V, Foti S, et al. Cysteine Oxidations in Mitochondrial Membrane Proteins: The Case of VDAC Isoforms in Mammals. Frontiers in cell and developmental biology. 2020;8:397. PubMed PMID: 32582695. Pubmed Central PMCID: 7287182.

21. Saletti R, Reina S, Pittala MGG, Magri A, Cunsolo V, Foti S, et al. Post-translational modifications of VDAC1 and VDAC2 cysteines from rat liver mitochondria. Biochimica et biophysica acta Bioenergetics. 2018 Sep;1859(9):806–16. PubMed PMID: 29890122.

22. Oliva M, De Pinto V, Barsanti P, Caggese C. A genetic analysis of the porin gene encoding a voltage-dependent anion channel protein in Drosophila melanogaster. Molecular genetics and genomics : MGG. 2002 Aug;267(6):746–56. PubMed PMID: 12207222.

23. Magri A, Di Rosa MC, Orlandi I, Guarino F, Reina S, Guarnaccia M, et al. Deletion of Voltage-Dependent Anion Channel 1 knocks mitochondria down triggering metabolic rewiring in yeast. Cellular and molecular life sciences : CMLS. 2019 Oct 26. PubMed PMID: 31655859.

24. Specchia V, Guarino F, Messina A, Bozzetti MP, De Pinto V. Porin isoform 2 has a different localization in Drosophila melanogaster ovaries than porin 1. Journal of bioenergetics and biomembranes. 2008 Jun;40(3):219–26. PubMed PMID: 18686020.

25. Hinsch KD, Asmarinah, Hinsch E, Konrad L. VDAC2 (porin-2) expression pattern and localization in the bovine testis. Biochimica et biophysica acta. 2001 Apr 16;1518(3):329–33. PubMed PMID: 11311949.

26. Pan L, Qiu D, Li J, Li J, Xu P, Zhao D, et al. Idiopathic male infertility in the Han population in China is affected by polymorphism in the VDAC2 gene. Oncotarget. 2016 Dec 13;7(50):82594–601. PubMed PMID: 27806320. Pubmed Central PMCID: 5347716.

27. Pan L, Liu Q, Li J, Wu W, Wang X, Zhao D, et al. Association of the VDAC3 gene polymorphism with sperm count in Han-Chinese population with idiopathic male infertility. Oncotarget. 2017 Jul 11;8(28):45242–8. PubMed PMID: 28431403. Pubmed Central PMCID: 5542182.

28. Sampson MJ, Lovell RS, Craigen WJ. The murine voltage-dependent anion channel gene family. Conserved structure and function. The Journal of biological chemistry. 1997 Jul 25;272(30):18966–73. PubMed PMID: 9228078.

29. Yuan J, Zhang Y, Sheng Y, Fu X, Cheng H, Zhou R. MYBL2 guides autophagy suppressor VDAC2 in the developing ovary to inhibit autophagy through a complex of VDAC2-BECN1-BCL2L1 in mammals. Autophagy. 2015;11(7):1081–98. PubMed PMID: 26060891. Pubmed Central PMCID: 4590641.

30. Xu A, Hua Y, Zhang J, Chen W, Zhao K, Xi W, et al. Abnormal Hypermethylation of the VDAC2 Promoter is a Potential Cause of Idiopathic Asthenospermia in Men. Scientific reports. 2016 Nov 28;6:37836. PubMed PMID: 27892527. Pubmed Central PMCID: 5124954.

31. Zhu Y, Liu Q, Liao M, Diao L, Wu T, Liao W, et al. Overexpression of lncRNA EPB41L4A-AS1 Induces Metabolic Reprogramming in Trophoblast Cells and Placenta Tissue of Miscarriage. Molecular therapy Nucleic acids. 2019 Dec 6;18:518–32. PubMed PMID: 31671345. Pubmed Central PMCID: 6838551.

32. Consortium GT. Human genomics. The Genotype-Tissue Expression (GTEx) pilot analysis: multitissue gene regulation in humans. Science. 2015 May 8;348(6235):648–60. PubMed PMID: 25954001. Pubmed Central PMCID: 4547484.

33. Noguchi S, Arakawa T, Fukuda S, Furuno M, Hasegawa A, Hori F, et al. FANTOM5 CAGE profiles of human and mouse samples. Scientific data. 2017 Aug 29;4:170112. PubMed PMID: 28850106. Pubmed Central PMCID: 5574368.

34. De Pinto V, Guarino F, Guarnera A, Messina A, Reina S, Tomasello FM, et al. Characterization of human VDAC isoforms: a peculiar function for VDAC3? Biochimica et biophysica acta. 2010 Jun-Jul;1797(6-7):1268–75. PubMed PMID: 20138821.

35. Deng W, Roberts SG. A core promoter element downstream of the TATA box that is recognized by TFIIB. Genes & development. 2005 Oct 15;19(20):2418–23. PubMed PMID: 16230532. Pubmed Central PMCID: 1257396.

36. Nepal C, Hadzhiev Y, Balwierz P, Tarifeno-Saldivia E, Cardenas R, Wragg JW, et al. Dual-initiation promoters with intertwined canonical and TCT/TOP transcription start sites diversify transcript processing. Nature communications. 2020 Jan 10;11(1):168. PubMed PMID: 31924754. Pubmed Central PMCID: 6954239.

37. Grandori C, Cowley SM, James LP, Eisenman RN. The Myc/Max/Mad network and the transcriptional control of cell behavior. Annual review of cell and developmental biology. 2000;16:653–99. PubMed PMID: 11031250.

38. Thiel G, Cibelli G. Regulation of life and death by the zinc finger transcription factor Egr-1. Journal of cellular physiology. 2002 Dec;193(3):287–92. PubMed PMID: 12384981.

39. Liu ZH, Dai XM, Du B. Hes1: a key role in stemness, metastasis and multidrug resistance. Cancer biology & therapy. 2015;16(3):353–9. PubMed PMID: 25781910. Pubmed Central PMCID: 4622741.

40. Triner D, Castillo C, Hakim JB, Xue X, Greenson JK, Nunez G, et al. Myc-Associated Zinc Finger Protein Regulates the Proinflammatory Response in Colitis and Colon Cancer via STAT3 Signaling. Molecular and cellular biology. 2018 Nov 15;38(22). PubMed PMID: 30181395. Pubmed Central PMCID: 6206459.

41. Woo AJ, Kim J, Xu J, Huang H, Cantor AB. Role of ZBP-89 in human globin gene regulation and erythroid differentiation. Blood. 2011 Sep 29;118(13):3684–93. PubMed PMID: 21828133. Pubmed Central PMCID: 3186340.

42. Qu H, Qu D, Chen F, Zhang Z, Liu B, Liu H. ZBTB7 overexpression contributes to malignancy in breast cancer. Cancer investigation. 2010 Jul;28(6):672–8. PubMed PMID: 20394500.

43. Niederreither K, Dolle P. Retinoic acid in development: towards an integrated view. Nature reviews Genetics. 2008 Jul;9(7):541–53. PubMed PMID: 18542081.

44. Matson JP, Cook JG. Cell cycle proliferation decisions: the impact of single cell analyses. The FEBS journal. 2017 Feb;284(3):362–75. PubMed PMID: 27634578. Pubmed Central PMCID: 5296213.

45. Allevato M, Bolotin E, Grossman M, Mane-Padros D, Sladek FM, Martinez E. Sequence-specific DNA binding by MYC/MAX to low-affinity non-E-box motifs. PloS one. 2017;12(7):e0180147. PubMed PMID: 28719624. Pubmed Central PMCID: 5515408.

46. Dong JT, Chen C. Essential role of KLF5 transcription factor in cell proliferation and differentiation and its implications for human diseases. Cellular and molecular life sciences : CMLS. 2009 Aug;66(16):2691–706. PubMed PMID: 19448973.

47. Scarpulla RC, Vega RB, Kelly DP. Transcriptional integration of mitochondrial biogenesis. Trends in endocrinology and metabolism: TEM. 2012 Sep;23(9):459–66. PubMed PMID: 22817841. Pubmed Central PMCID: 3580164.

48. Hodneland Nilsson LI, Nitschke Pettersen IK, Nikolaisen J, Micklem D, Avsnes Dale H, Vatne Rosland G, et al. A new live-cell reporter strategy to simultaneously monitor mitochondrial biogenesis and morphology. Scientific reports. 2015 Nov 24;5:17217. PubMed PMID: 26596249. Pubmed Central PMCID: 4657046.

49. Costoya JA. Functional analysis of the role of POK transcriptional repressors. Briefings in functional genomics & proteomics. 2007 Mar;6(1):8–18. PubMed PMID: 17384421.

50. Labrecque MP, Prefontaine GG, Beischlag TV. The aryl hydrocarbon receptor nuclear translocator (ARNT) family of proteins: transcriptional modifiers with multi-functional protein interfaces. Current molecular medicine. 2013 Aug;13(7):1047–65. PubMed PMID: 23116263.

51. Hayes JD, Dinkova-Kostova AT. The Nrf2 regulatory network provides an interface between redox and intermediary metabolism. Trends in biochemical sciences. 2014 Apr;39(4):199–218. PubMed PMID: 24647116.

52. Anckar J, Sistonen L. Regulation of HSF1 function in the heat stress response: implications in aging and disease. Annual review of biochemistry. 2011;80:1089–115. PubMed PMID: 21417720.

53. Morgan R, Pandha HS. PBX3 in Cancer. Cancers. 2020 Feb 13;12(2). PubMed PMID: 32069812. Pubmed Central PMCID: 7072649.

54. Saghizadeh M, Akhmedov NB, Yamashita CK, Gribanova Y, Theendakara V, Mendoza E, et al. ZBED4, a BED-type zinc-finger protein in the cones of the human retina. Investigative ophthalmology & visual science. 2009 Aug;50(8):3580–8. PubMed PMID: 19369242. Pubmed Central PMCID: 2848067.

55. Oh S, Hwang ES. The role of protein modifications of T-bet in cytokine production and differentiation of T helper cells. Journal of immunology research. 2014;2014:589672. PubMed PMID: 24901011. Pubmed Central PMCID: 4036734.

56. Li N, Zhao CT, Wang Y, Yuan XB. The transcription factor Cux1 regulates dendritic morphology of cortical pyramidal neurons. PloS one. 2010 May 11;5(5):e10596. PubMed PMID: 20485671. Pubmed Central PMCID: 2868054.

57. Hidaka K, Yamamoto I, Arai Y, Mukai T. The MEF-3 motif is required for MEF-2-mediated skeletal muscle-specific induction of the rat aldolase A gene. Molecular and cellular biology. 1993 Oct;13(10):6469–78. PubMed PMID: 8413246. Pubmed Central PMCID: 364706.

58. Kageyama R, Shimojo H, Ohtsuka T. Dynamic control of neural stem cells by bHLH factors. Neuroscience research. 2019 Jan;138:12–8. PubMed PMID: 30227160.

59. Buratowski S, Zhou H. Functional domains of transcription factor TFIIB. Proceedings of the National Academy of Sciences of the United States of America. 1993 Jun 15;90(12):5633–7. PubMed PMID: 8516312. Pubmed Central PMCID: 46775.

60. Fang X, Huang Z, Zhou W, Wu Q, Sloan AE, Ouyang G, et al. The zinc finger transcription factor ZFX is required for maintaining the tumorigenic potential of glioblastoma stem cells. Stem cells. 2014 Aug;32(8):2033–47. PubMed PMID: 24831540. Pubmed Central PMCID: 4349564.

61. Basso K, Dalla-Favera R. BCL6: master regulator of the germinal center reaction and key oncogene in B cell lymphomagenesis. Advances in immunology. 2010;105:193–210. PubMed PMID: 20510734.

62. Bai YQ, Miyake S, Iwai T, Yuasa Y. CDX2, a homeobox transcription factor, upregulates transcription of the p21/WAF1/CIP1 gene. Oncogene. 2003 Sep 11;22(39):7942–9. PubMed PMID: 12970742.

63. Friedman JR, Kaestner KH. The Foxa family of transcription factors in development and metabolism. Cellular and molecular life sciences : CMLS. 2006 Oct;63(19-20):2317–28. PubMed PMID: 16909212.

64. Suzuki H, Ahn HW, Chu T, Bowden W, Gassei K, Orwig K, et al. SOHLH1 and SOHLH2 coordinate spermatogonial differentiation. Developmental biology. 2012 Jan 15;361(2):301–12. PubMed PMID: 22056784. Pubmed Central PMCID: 3249242.

65. He S, Yang S, Niu M, Zhong Y, Dan G, Zhang Y, et al. HMG-box transcription factor 1: a positive regulator of the G1/S transition through the Cyclin-CDK-CDKI molecular network in nasopharyngeal carcinoma. Cell death & disease. 2018 Jan 24;9(2):100. PubMed PMID: 29367693. Pubmed Central PMCID: 5833394.

66. Wang W, Lufkin T. Hmx homeobox gene function in inner ear and nervous system cell-type specification and development. Experimental cell research. 2005 Jun 10;306(2):373–9. PubMed PMID: 15925593.

67. Mahnke J, Schumacher V, Ahrens S, Kading N, Feldhoff LM, Huber M, et al. Interferon Regulatory Factor 4 controls TH1 cell effector function and metabolism. Scientific reports. 2016 Oct 20;6:35521. PubMed PMID: 27762344. Pubmed Central PMCID: 5071867.

68. Jennings W, Hou M, Perterson D, Missiuna P, Thabane L, Tarnopolsky M, et al. Paraspinal muscle ladybird homeobox 1 (LBX1) in adolescent idiopathic scoliosis: a cross-sectional study. The spine journal : official journal of the North American Spine Society. 2019 Dec;19(12):1911–6. PubMed PMID: 31202838.

69. Musa J, Aynaud MM, Mirabeau O, Delattre O, Grunewald TG. MYBL2 (B-Myb): a central regulator of cell proliferation, cell survival and differentiation involved in tumorigenesis. Cell death & disease. 2017 Jun 22;8(6):e2895. PubMed PMID: 28640249. Pubmed Central PMCID: 5520903.

70. Zhao M, Mishra L, Deng CX. The role of TGF-beta/SMAD4 signaling in cancer. International journal of biological sciences. 2018;14(2):111–23. PubMed PMID: 29483830. Pubmed Central PMCID: 5821033.

71. Moore BD, Jin RU, Lo H, Jung M, Wang H, Battle MA, et al. Transcriptional Regulation of X-Box-binding Protein One (XBP1) by Hepatocyte Nuclear Factor 4alpha (HNF4Alpha) Is Vital to Beta-cell Function. The Journal of biological chemistry. 2016 Mar 18;291(12):6146–57. PubMed PMID: 26792861. Pubmed Central PMCID: 4813565.

72. Wang F, Qiang Y, Zhu L, Jiang Y, Wang Y, Shao X, et al. MicroRNA-7 downregulates the oncogene VDAC1 to influence hepatocellular carcinoma proliferation and metastasis. Tumour biology : the journal of the International Society for Oncodevelopmental Biology and Medicine. 2016 Aug;37(8):10235–46. PubMed PMID: 26831666.

73. Cam H, Balciunaite E, Blais A, Spektor A, Scarpulla RC, Young R, et al. A common set of gene regulatory networks links metabolism and growth inhibition. Molecular cell. 2004 Nov 5;16(3):399–411. PubMed PMID: 15525513.

74. Hughes TR. A handbook of transcription factors. .Springer Science+Business Media B.V. 2011(Subcellular Biochemistry 52).

75. Guarino FZ, F.; Mela, L.; Pappalardo, X.; Messina, A.; De Pinto, V. NRF-1 and HIF-1α modulate activity of human VDAC1 gene promoter during starvation and hypoxia in HeLa cells. Biorxiv. 2020.

76. Camara AKS, Zhou Y, Wen PC, Tajkhorshid E, Kwok WM. Mitochondrial VDAC1: A Key Gatekeeper as Potential Therapeutic Target. Frontiers in physiology. 2017;8:460. PubMed PMID: 28713289. Pubmed Central PMCID: 5491678.

77. Cartharius K, Frech K, Grote K, Klocke B, Haltmeier M, Klingenhoff A, et al. MatInspector and beyond: promoter analysis based on transcription factor binding sites. Bioinformatics. 2005 Jul 1;21(13):2933–42. PubMed PMID: 15860560.

78. Fornes O, Castro-Mondragon JA, Khan A, van der Lee R, Zhang X, Richmond PA, et al. JASPAR 2020: update of the open-access database of transcription factor binding profiles. Nucleic acids research. 2020 Jan 8;48(D1):D87–D92. PubMed PMID: 31701148. Pubmed Central PMCID: 7145627.

79. Gheorghe M, Sandve GK, Khan A, Cheneby J, Ballester B, Mathelier A. A map of direct TF-DNA interactions in the human genome. Nucleic acids research. 2019 Feb 28;47(4):e21. PubMed PMID: 30517703. Pubmed Central PMCID: 6393237.

